# Maintaining Gene Expression Levels by Positive Feedback in Burst Size in the Presence of Infinitesimal Delay

**DOI:** 10.1101/379602

**Authors:** Pavol Bokes

**Affiliations:** Department of Applied Mathematics and Statistics, Comenius University, Bratislava 842 48, Slovakia

## Abstract

Synthesis of individual molecules in the expression of genes often occurs in bursts of multiple copies. Gene regulatory feedback can affect the frequency with which these bursts occur or their size. Whereas frequency regulation has traditionally received more attention, we focus specifically on the regulation of burst size. It turns out that there are (at least) two alternative formulations of feedback in burst size. In the first, newly produced molecules immediately partake in feedback, even within the same burst. In the second, there is no within-burst regulation due to what we call infinitesimal delay. We describe both alternatives using a minimalistic Markovian drift-jump framework combining discrete and continuous dynamics. We derive detailed analytic results and efficient simulation algorithms for positive noncooperative autoregulation (whether infinitesimally delayed or not). We show that at steady state both alternatives lead to a gamma distribution of protein level. The steady-state distribution becomes available only after a transcritical bifurcation point is passed. Interestingly, the onset of the bifurcation is postponed by the inclusion of infinitesimal delay.

## 1. Introduction

Stochastic expression of individual genes drives random fluctuations in the concentration of protein molecules in individual cells and inhomogeneity of protein concentration across cell populations [1, 2]. The production of protein molecules in bursts, i.e. batches of many molecules within brief periods of time, has been identified as a major source of gene-expression variability [3, 4].

The presence of feedback of a protein on its own gene expression is ubiquitous in biological circuits [5]. If protein is produced in bursts, regulatory feedback can act in two fundamentally different ways: it can affect the frequency with which bursts occur in time [6]; alternatively, it can affect the size of the bursts [7]. Feedback can be negative and positive; here, we shall specifically focus on positive feedback on burst size. The impact of negative feedback on burst size and burst frequency has been investigated in [8, 9]

Models of stochastic gene expression are typically based on the random telegraph framework [10–17]. In a random telegraph model, a gene can be either in an On state or in an Off state, transitioning randomly in time between the two. In addition to the binary variable indicating whether the gene is On and Off, there is another time-dependent numerical variable indicating the amount of protein in the cell. In many models the protein amount is considered to be a discrete variable [18–21], but here we use a hybrid discrete-continuous formalism [22–29] and treat protein level as a continuous variable; we refer to it also as concentration. The protein concentration increases whenever the gene is On and decreases when the gene is Off.

The mathematical abstraction of On and Off states can have alternative biological interpretations in different contexts. They can represent active and inactive states of the promoter region of the gene [30]. Alternatively, they can indicate the presence or absence of a messenger transcript molecule [31].

The random telegraph model can generate bursts if the time spent in the On state is short but accompanied by rapid protein synthesis (Figures 1A). Bursts lead to rapid but continuous increase in the protein concentration occurring on a fast timescale (Figure 1B). In the singular limit of infinitesimally brief On periods, the random telegraph model reduces to a jump-drift model for protein level (Figure 1C). The state of the gene is no longer explicitly needed in the reduced model; burst are represented by discontinuous jumps in protein concentration (Figure 1C).

**Figure 1.**
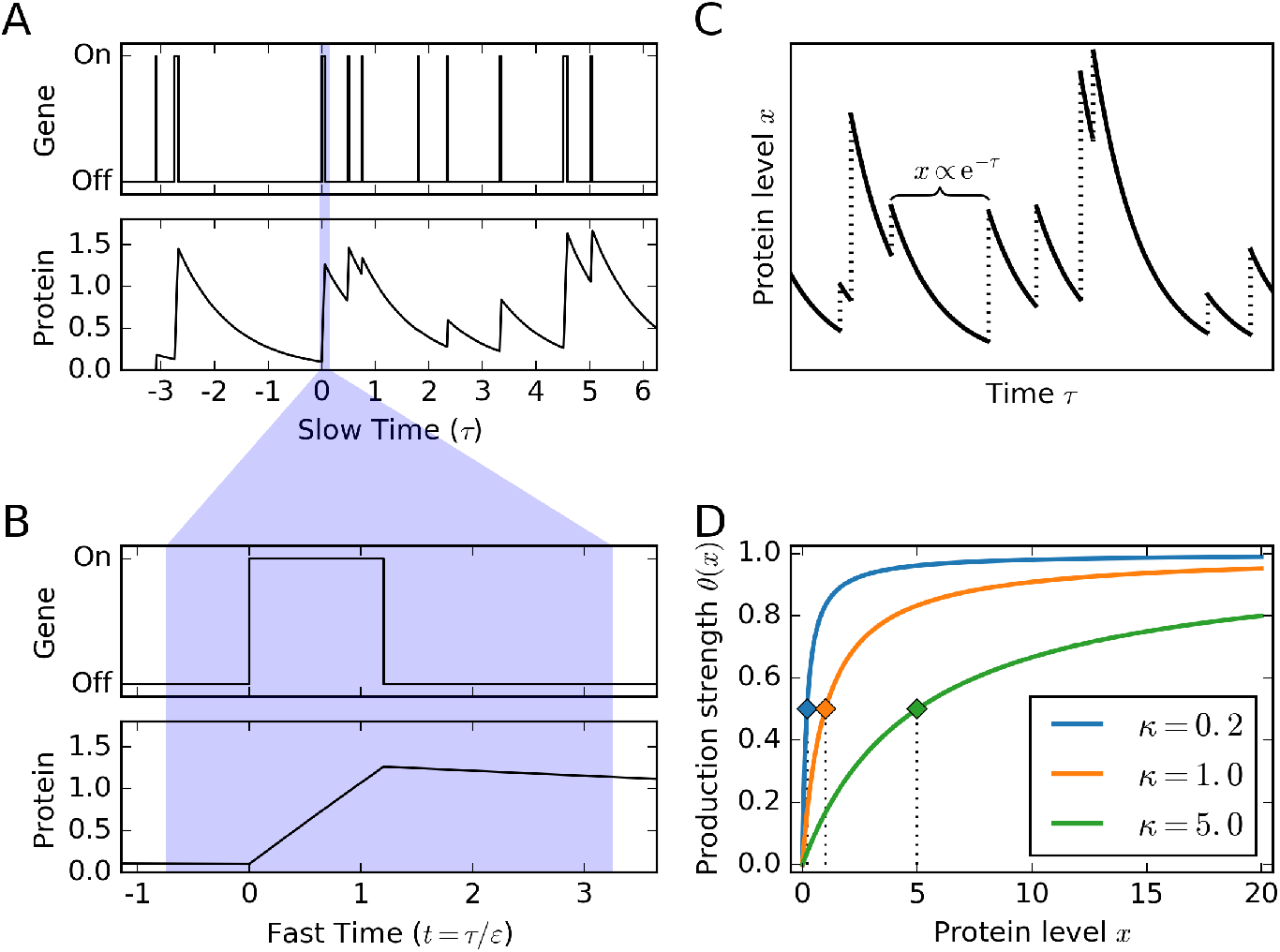
*A*: Random telegraph model for gene expression. Protein concentration increases if gene is On and decreases if gene is Off. Short On periods accompanied by fast production lead to burst-like gene expression. *B*: The burst dynamics can be deciphered by looking on the fast timescale (the scaling factor is set to ε = 0.05 here). *C*: Protein dynamics in the reduced drift-jump model. Between bursts (vertical dotted lines) the protein level decays exponentially (solid lines; one such period is demarcated by the braces). *D*: Positive feedback response function of the Michaelis-Menten type (11) as function of protein level *x* for selected values of the threshold level *κ*. The dotted black lines ending in coloured markers indicate that the response is half-maximal at *x* = *κ*.

We will perform the bulk of our analysis on the reduced model with discontinuous bursts. Nevertheless, the full random-telegraph model will initially be used to derive the burst-size distribution. Importantly, the analysis of the random-telegraph model will imply that there are two fundamentally different ways of implementing feedback in burst size. First, feedback can act immediately at the level of fast timescale of burst growth. Alternatively, feedback can only react on the changes on the slow timescale. The latter differs from the former in the inclusion of what we call an infinitesimal delay. It has been shown that that the inclusion of an infinitesimal delay in negative feedback on burst size can potentially destabilise the noise-controlling capabilities of the negative feedback loop [9].

The purpose of the present work is to examine the effects of positive feedback on burst size. For specificity, we consider non-cooperative Michaelis-Menten regulation (Figure 1D). We investigate the effect of the feedback on the onset of a transcritical bifurcation in the model. We focus systematically on the differences in the two versions of the model which differ in the omission/inclusion of the infinitesimal delay. The burst-size distributions are derived in Section 2. The master equations are formulated and solved in Section 3. Exact stochastic simulation algorithms are formulated in Section 4 and used in Section 5 to cross-validate analytic results. The paper ends with a summary and discussion in Section 6.

## 2. Burst dynamics

Here we use the random telegraph model (Figure 1A) to examine the consequences of the fast-timescale burst dynamics (Figure 1B). The deliverable is to derive the burst-size distributions, which will serve as building blocks of jump-drift models in later Sections. Two versions of the model are treated separately: one in which feedback acts immediately; the other in which feedback operates with an infinitesimal delay.

### 2.1. Undelayed case

Without loss of generality we assume that a burst starts at time *t* = 0 and that its duration *T* is exponentially distributed, i.e.

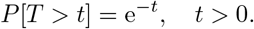

We also assume that within the burst duration the protein level grows according to an ordinary differential equation

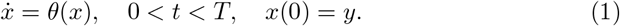

The value *y* gives the before-burst protein level. The function *θ*(*x*), which is assumed to be positive, characterises the feedback response. Later we will restrict ourselves to a specific non-cooperative Michaelis-Menten-type form (Figure 1D). The after-burst protein level is a random variable which is defined by

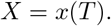

By the burst kernel *B*_1_(*x*|*y*) we will understand the right-tail cumulative distribution function of *X*. It follows that the burst kernel is given by

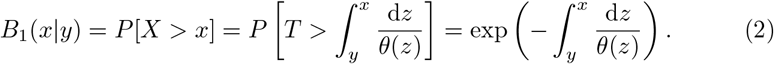

The subscript one in the burst kernel indicates that this is the first formulation of the burst-size feedback. The second one follows.

### 2.2. Infinitesimally delayed case

We modify the differential equation into

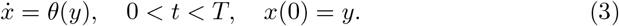

Note that in (3) the growth rate is constant within the burst and depends only on the initial before-burst protein level. The burst kernel *B*_2_(*x*|*y*) is now given by

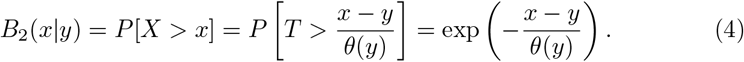

The burst kernels (2) and (4) will be used in the jump-drift master equation in the next section. The bursting timescale of this section will thereby be assumed to be infinitesimally small. Therefore we refer to the delay that distinguishes (4) from (2) as infinitesimal delay.

## 3. Master equation

Here we build a model that characterises the slow-timescale behaviour of the reduced jump-drift model (Figure 1C). The burst kernels derived in Section 2 considering the fast timescale will serve as building blocks. It is assumed that bursts occur with rate α and that between bursts the protein concentration decays exponentially. Without loss of generality one can assume that the decay rate constant is equal to one. Under such assumptions, the probability *p*(*x, τ*) of observing the protein at level *x* at time *τ* satisfies a partial integro-differential equation

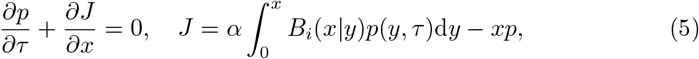

where *B_i_*(*x*|*y*) is either one of the kernels (2) or (4) depending on whether feedback is immediate or operates with infinitesimal delay. The parameter *α* gives the (constant) burst rate or burst frequency. The first equation in (5) is a differential form of probability conservation principle. The probability flux *J* is a sum of a nonlocal integral flux due to bursts and a local flux −*xp* due to the deterministic drift of protein concentration between bursts. Note that the time variable *τ* in (5) is much slower than the time variable *t* used in Section 2. More precisely, we may say that d*τ*/d*t* = *ε*, where *ε* ≪ 1 is an infinitesimally small timescale ratio (cf. Figure 1A,B).

In the absence of feedback, *θ*(*y*) ≡ *θ* is independent of the protein level, and *B_i_*(*x*|*y*) = exp(−(*x* − *y*)/*θ*). Equation (5) then reduces to a well known model for constitutive bursty protein synthesis [6]. In particular, it is known that the stationary distribution in the unregulated model is the gamma distribution with shape parameter α and scale parameter *θ* [6]. Below, we intend to extend this result to the case of positive feedback in burst size.

Equating the probability flux to zero, we obtain an integral equation

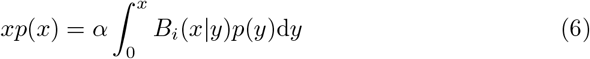

for the steady-state protein distribution. The rest of this chapter is devoted to solving this integral equation separately for *i* = 1 and *i* = 2. For its central position in this paper, we will refer to (6) (and its specifications) as the master equation. The master equation (6) is a singular homogeneous Volterra equation of the second kind [32]. It is shown below that the integral kernel *B*_1_(*x*|*y*) allows for an explicit solution. On the other hand, the integral kernel *B*_2_(*x*|*y*) leads to tractable equations only in specific situations.

### 3.1. Undelayed case

(*i* = 1). The kernel (2) can be factorised into

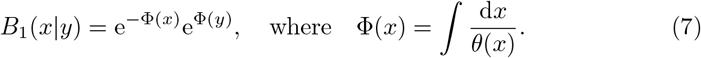

Integral equations with factorised kernels can be transformed into a differential equation [32]. Indeed, inserting (7) into (6) yields

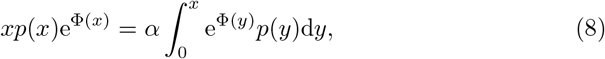

and differentiation gives

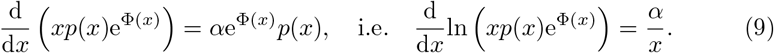

Integrating (9) yields

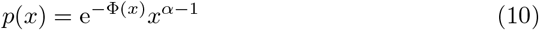

up to a normalisation constant. Below we will focus specifically on a Michaelis-Menten type regulation

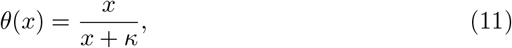

where *κ* represents a critical concentration that is required to achieve half-maximal production (cf. Figure 1D). The maximal production can be scaled to one without loss of generality.

For Michaelis-Menten type regulation we easily find that

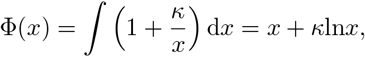

so that the steady-state solution (10) simplifies to

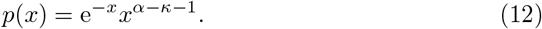

If *α* > *κ*, then (12) can be normalised into the gamma probability distribution with shape *α* − *κ* and scale one. The steady state mean is then given by the product of shape and scale which is also α — κ. If on the other hand *α* < *κ*, the master equation does not possess normalised solutions, and the protein concentration will decrease to zero with probability one. In summary, the protein mean is given by

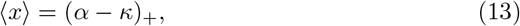

where (*a*)_+_ = max(*a*, 0) is the positive part of a real number *a*. Interestingly, the steady-state mean of the undelayed stochastic formulation coincides with the global attractor of the deterministic formulation

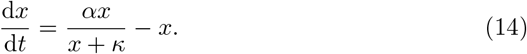

Differential equation (14) exhibits a transcritical bifurcation at *κ* = *α* [33]. There are two steady states, 0 and *α* − *κ*, which coalesce at the point of bifurcation and exchange stability. Consequently, the global attractor of (14) is given by (*α* − *κ*)_+_, the same value as the steady-state mean (13) of the undelayed stochastic formulation. Below we examine how the inclusion of infinitesimal delay affects the bifurcation behaviour in the stochastic model.

### 3.2. Infinitesimally delayed case

(*i* = 2). Unlike the undelayed burst kernel (2), the burst kernel (4) is not factorisable. A general solution formula for the master equation (6) with burst kernel (4) is unavailable. However, progress can be made if the regulatory function is of Michaelis-Menten type (11), in which case the master equation (6) takes the specific form of

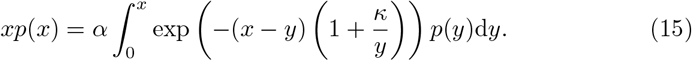

Multiplying both sides of 15 by e^*x*^ simplifies it into

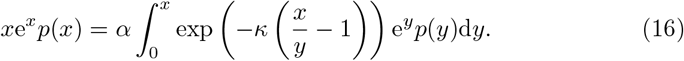

We look for a solution in the form

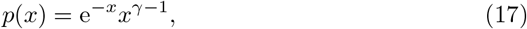

where *γ* is a real constant yet to be determined. Inserting (17) into (16) we see that

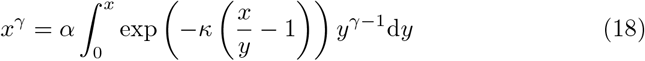

is required to hold for all *x* > 0. Substitution *x*/*y* − 1 = ξ in the integral on the right-hand side yields

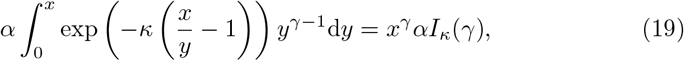

where

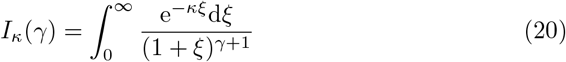

is a special function dependent on *γ* but independent of *x*. Equating the left-hand side of (18) to the right-hand side of (19) and dividing the result by *x^γ^* we find that

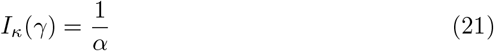

is a necessary and sufficient condition for (17) to solve the master equation (6).

The function *I_κ_*(*γ*) is decreasing, and maps the interval (−∞, ∞) onto the interval (0, ∞). Therefore for a given *α* > 0 there exists a unique solution γ to the algebraic equation (21) which can formally be written as

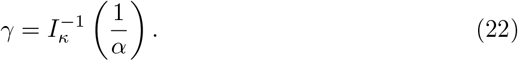

The function *γ* = *γ_κ_*(*α*) defined through the formula (22) is an increasing function which maps the interval (0, ∞) onto (−∞, ∞). It crosses zero at a value *α*_bif_ = 1/*I_κ_*(0). The solution (17), where *γ* is given by (22), can be normalised into a probability density function only after α passes the bifurcation value *α*_bif_. The condition *α* > *α*_bif_ represents the analogue of the condition *α* > *κ* required for the solution (12) to the undelayed problem to be integrable. Prior to the bifurcation point, i.e. for *α* ≤ *α*_bif_, the random process giving the temporal dynamics of protein concentration decreases to zero with probability one. It follows that the protein mean is given by

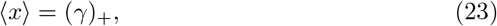

which represents an infinitesimally delayed analogue of (13).

Below, we derive inequalities that help establish bounds on the protein mean (23) in the infinitesimally delayed model. For *γ* ≥ 0 the integrand in (20) can be bounded from below and above by

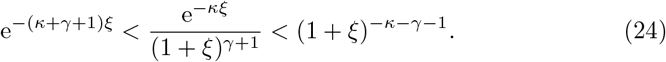

Integrating (24) with respect to *ξ* from zero to infinity, and using (20) and (21), yields

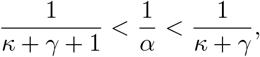

i.e.

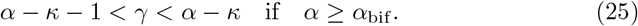

The second inequality (25) implies that the protein mean (23) in the infinitesimally delayed model is strictly smaller than the mean (13) in the undelayed model. Specifically for *α* = *α*_bif_, for which *γ* = 0, the second inequality (25) reduces to *α*_bif_ > *κ*. Recall that *α* = *κ* gives the bifurcation condition in the undelayed and deterministic models. Therefore, we can conclude that the introduction of infinitesimal delay postpones the onset of the transcritical bifurcation.

The first inequality in (25) provides an upper bound on the bifurcation postponement. In particular, it implies that *α*_bif_ < *κ*+1, i.e. that the onset of the bifurcation cannot be delayed by more than one in the chosen units of burst frequency.

## 4. Exact stochastic simulation algorithm

The stochastic simulation algorithm iterates three essential steps: the drawing of a waiting time until the upcoming burst; the discounting of the protein level during the waiting period owing to protein decay; and the increase of the protein level by a randomly sampled burst size. The first and second steps are the same for both models (the undelayed and the infinitesimally delayed) of burst-size regulation. The models differ in the third step of the algorithm. Let us start with the common steps.

Neither model incorporates feedback in burst frequency. Therefore, the burst occurrences form a Poisson process whose intensity is given by the burst frequency parameter *α*. The waiting times between successive bursts are exponentially distributed with mean 1/*α*. Consequently, the waiting time *τ_i_* between the (*i* − 1)-th and *i*-th bursts can be drawn using the well-known formula for drawing exponentially distributed variates,

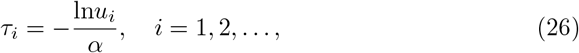

in which *u_i_, i* = 1,… are random variates drawn independently of each other from the unit-interval uniform distribution.

Denoting by *y_i_* and *x_i_* the protein level immediately before and immediately after the i-th burst, the second step of the simulation rests in discounting the amount of protein degraded during the period between successive bursts,

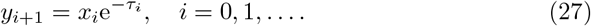

Applying formula (27) for *i* = 0, we obtain the protein level *y*_1_ immediately before the first burst from an initial value *x*_0_, which is specified in the input of the algorithm.

We are now in position to unveil the third and final step, which samples the post-burst level *x_i_* based on the knowledge of the pre-burst level *y_i_*. As advertised above, the step takes a different form depending on the choice of the model. In the infinitesimally delayed case, we will obtain a straightforward implementation which can be universally applied regardless of the choice of the feedback response functions *θ*(*y*). In the undelayed case, the implementation is more delicate and hinges on the specifics of the chosen Michaelis-Menten-type regulation (11). Methodologically, this represents a reversal of the situation encountered in Section 3, where the undelayed model was shown to admit an explicit steady-state distribution for any choice of *θ*(*y*), whereas in the infinitesimally delayed case an explicit solution was found only for the specific Michaelis-Menten regulation. Let us proceed with the description of the post-burst level sampling procedure in the simpler case, here being that of the infinitesimally delayed model.

### 4.1. Infinitesimally delayed model

The burst-size distribution (4), conditioned on knowing the pre-burst level *y*, is exponentially distributed with mean *θ*(*y*). The level *x_i_* of protein after the *i*-th burst can therefore be sampled by

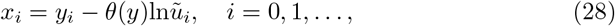

where *y_i_* is the corresponding pre-burst level and *ũ_i_* are random variates drawn from the unit-interval uniform distribution independently of each other and of any other random variates.

### 4.2. Undelayed model

In the Michaelis-Menten type regulation (11), the cumulative distribution function (2) of the post-burst level *X* satisfies

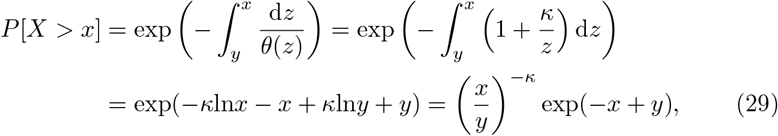

where *y* is the pre-burst level and *x* > *y* is any admissible post-burst level. Next we discuss how post-protein level *x* can be sampled from the distribution (29). One option would be to use the inversion sampling method directly on (29). This would involve equating the right-hand side of (29) to a randomly drawn variate from the unit interval and solving in terms of the post-protein level *x*. However, since this means that a transcendental equation needs to be solved, we choose to use a different sampling procedure.

We define two independent auxiliary variables *X*^(1)^ and *X*^(2)^ with distributions

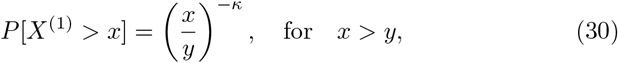

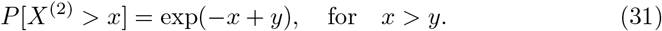

The minimum of these two has distribution

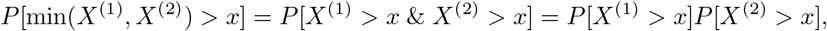

which coincides with the distribution (29) of the post-burst level. The post-burst level can therefore be sampled by drawing random variates *x*^(1)^ and *x*^(2)^ from the distributions (30) and (31) and then taking the minimal of the two. Drawing from the distributions (30) and (31) can readily be done using the inversion sampling method.

Denoting, as before, the *i*-th burst’s pre- and post-levels by *y_i_* and *x_i_*, we find that

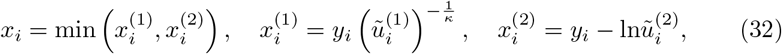

in which 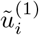 and 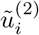 are random variates drawn from the uniform distribution on the unit interval independently of each other and of any previous ones.

## 5. Cross-validation of analysis with simulations

Adapting a method previously used in [9], we estimate the steady-state protein mean 〈*x*〉 by the time average 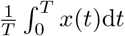, where *T* ≫ 1, of a sample trajectory *x*(*t*). The trajectory decays exponentially between successive bursts, so that

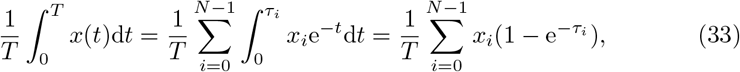

where *y_i_* and *x_i_* are the pre- and post-burst protein levels, *τ_i_* give the waiting times, and 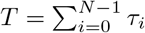, where *N* is a large integer. Given that *τ_i_* are independent and identically distributed with mean 1/*α*, we have *T* ≈ *N*/*α* by the law of large numbers.

Numerical experiments suggest that the temporal average (33) can be sensitive to the choice of the initial value *x*(0) = *x*_0_, which is typically arbitrarily chosen (we use *x*_0_ = 1 throughout). In order to reduce the influence of the initial data, we can shift the lower bound in the sum (33) from zero to a large positive integer *M*; to retain the number of summands we can move the upper bound from *N* to *N* + *M* − 1. This leads to

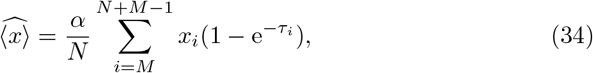

where we also replaced 1/*T* by its law-of-large-numbers estimate *α*/*N*. We use *N* = 10^5^ and *M* = 10^4^ throughout. The sequences *y_i_, x_i_*, and *τ_i_* are obtained iteratively from (26), (27), and (28) in the infinitesimally delayed case or from (32) in the undelayed case. In order to obtain additional ensemble averaging, we perform a hundred independent simulations to obtain a hundred independent estimates (34) of protein mean. Each reported value of the protein mean is equal to the average over the hundred estimates. We also calculate the standard deviation of these estimates and the confidence intervals for the average value. In our simulations, the confidence intervals are extremely narrow, suggesting that we obtained very precise estimates of the actual steady-state protein mean.

Figure 2 shows the dependence of the steady-state protein mean 〈*x*〉 on the bifurcation parameter *α* (the burst frequency) for two selected values of the critical concentration *κ* = 0.2 (Fig. 2, left panel), and *κ* = 5.0 (Fig. 2, right panel). The theoretical results for the undelayed model (13) (orange lines) and for the infinitesimally delayed model (23) (blue lines) are cross-validated with the kinetic Monte Carlo estimate (34) (discrete markers). We observe an excellent agreement between the two. In both panels, the onset of the transcritical bifurcation is postponed in the infinitesimally delayed model. In the left panel of Fig. 2, the bifurcation value is *α*_bif_ = 0.69 in the infinitesimally delayed model and by *κ* = 0.2 in the undelayed model. In the right panel of Fig. 2, the bifurcation occurs at α_bi_f = 5.87 if the infinitesimal delay is included and at *κ* = 5 if it is omitted. Both panels demonstrate that after the bifurcation the protein mean *γ* of the infinitesimally delayed model stays below the mean *α* − *κ* of the undelayed model; this confirms the upper-bound result in (25). For large values of *κ* (such as *κ* = 5 in the right panel and larger), the mean *γ* of the infinitesimally delayed model is close to the lower-bound in (25).

**Figure 2.**
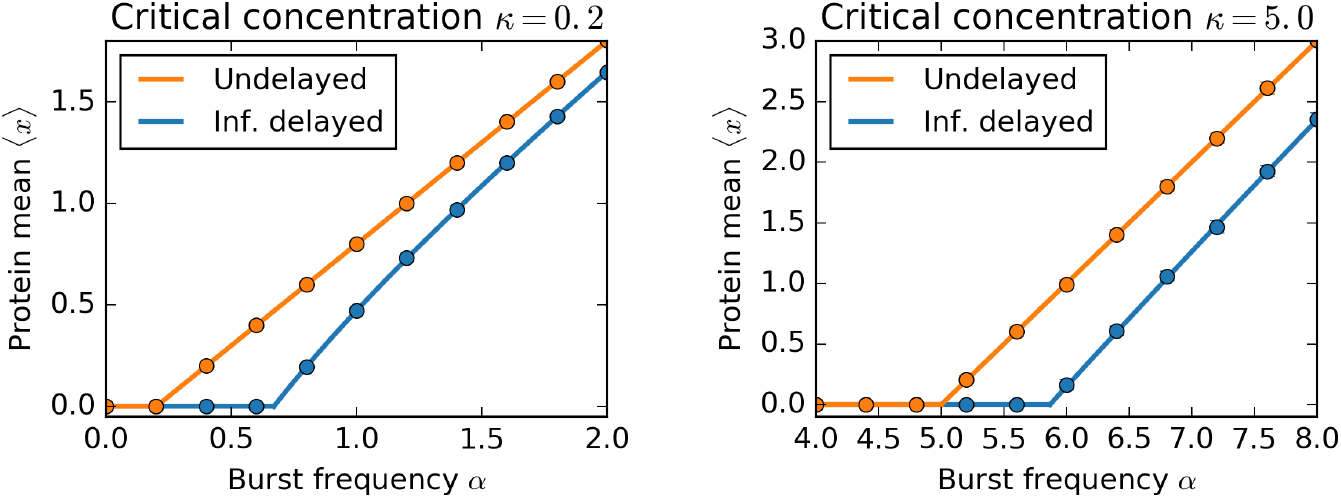
Transcritical bifurcation of steady-state protein mean in the undelayed and infinitesimally delayed models. Analytical results (13) and (23) (solid lines) are cross-validated with kinetic Monte Carlo estimates (34) (discrete markers). Each individual marker gives the average value based on one hundred independent estimates (34).

The positive branch of the solid blue line in Fig. 2 (either panel) is the graph of the function *γ* = *γ_κ_*(*α*). Its definition (22) involves taking the inverse of the special function *I_κ_*(*γ*) defined through a parametric integral (20). Nevertheless, for the purposes of plotting a graph it is not necessary to take the inverse, since it and the original function share the same graph. The parametric integral (20) can be expressed in terms of the confluent hypergeometric function of the second kind [34]

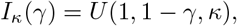

which is widely available in common mathematical software; we used the implementation in Python’s scientific computation library SciPy (version 0.17.0) which is accessible as `hyperu` from the module `scipy.special`.

## 6. Discussion

We investigated a stochastic model for the temporal dynamics of protein concentration x that positively autoregulates the size of its production bursts. Two versions of the model were considered, differing in the inclusion or omission of an infinitesimal delay in the feedback. The presence of infinitesimal delay means that on the fast timescale of individual bursts, newly produced protein do not immediately promote protein synthesis within the same burst. The effect of the protein concentration on the protein synthesis is characterised by the feedback response function *θ*(*x*). Some of our results apply to any choice of *θ*(*x*), whereas others are restricted to a Michaelis-Menten type form of the function. The two versions of the model differ in the burst kernels, Eqns (2) and (4), which give the probability of a burst to surpass concentration *x*, having started at concentration *y*.

The large-time behaviour of the model is characterised by the stationary distribution of protein concentration. We derived a Volterra integral equation for the stationary distribution. The bursting kernel appears as the integral kernel of the integral operator in the equation. For the undelayed model, the kernel is separable, and the integral equation admits an explicit solution regardless of the choice of response function *θ*(*x*). For the infinitesimally delayed model, we were able to determine the stationary distribution only for the Michaelis-Menten type response function. In both versions of the model, the stationary distribution is a gamma distribution for Michaelis-Menten type response function. As is well known, gamma distributions are a two-parametric family, one parameter giving the scale and the other the shape of the distribution. The scale of the stationary distribution can be set to one by a suitable choice of concentration units. The shape of the stationary distribution, which coincides with its mean, is a function of the dimensionless model parameters. For the undelayed model, the mean is given by an explicit function (13) of the parameters, which coincides with the attractor of an associated deterministic model (14). For the delayed model, on the other, the mean is given implicitly through an equation (21) involving a special function (20).

In addition to analytic results, we presented a simulational approach to the both versions of our model. The crucial step in the algorithm is to draw the post-burst protein concentration from the burst-size distribution given by the burst kernel. For the infinitesimally delayed model, the burst-size distribution is exponentially distributed, with the response function determining the dependence of the mean burst size on the pre-burst level. Since drawing from an exponential distribution is easily done using inversion sampling method, we obtain an exact and efficient algorithm for simulating sample paths in the infinitesimally delayed version of our model. For the undelayed version of our model, the burst-size distribution can assume nontrivial (and non-parametric) forms depending on the choice of the response function. In general, finding the post-burst protein level would require the use of numerical integration or some form of (possibly inefficient) rejection sampling method. However, restricting ourselves to the Michaelis-Menten type response function, we obtained an exact and efficient simulation algorithm that generates the post-burst level based on two samples from the uniform distribution on the unit interval.

We crossvalidated our analytic results with stochastic simulations in Section 5. Our results pertain to the onset of transcritical bifurcation in the positively autoregulated model. Transcritical bifurcation in deterministic dynamical systems occurs when two steady states collide and exchange stability in response to parametric change. The deterministic formulation of our model (14) presents a typical situation. Prior to the bifurcation, the trivial zero steady state is stable and a nontrivial steady state is unstable and unrealistic (negative). As production rate increases, a transcritical bifurcation occurs at which the two steady states coalesce at zero and exchange stability. After the bifurcation, the trivial zero steady state is unstable and the nontrivial steady state is positive, stable, and attracts all positive initial conditions.

An analogous behaviour is exhibited by both versions of our stochastic model. For simplicity, we refer to it also as transcritical bifurcation. There are two solutions to the master equation: a trivial one — the delta function giving all mass to zero concentration —, and a non-trivial one (22). Prior to the bifurcation, the non-trivial solution is inadmissible as it has an infinite *L*^1^ norm and cannot be normalised into a probability density function. Initial distributions tend to the trivial steady state. The bifurcation occurs as the parameter giving the burst frequency increases. After the bifurcation, the non-trivial solution can be normalised into the probability density function of a gamma distribution. It serves as the large-time attractor of all initial distributions. Our results imply that the inclusion of the infinitesimal delay postpones the onset of the transcritical bifurcation in the stochastic model in the sense that greater burst frequencies are required to obtain a self-sustainable steady-state distribution in an infinitesimally delayed model. After the bifurcation, the mean protein concentration in the infinitesimally delayed model remains lower than in the undelayed model (the latter being equal to the deterministic steady-state level).

In summary, we have investigated two versions of a stochastic model for pulsatile protein dynamics with feedback in burst size. Our methods combined differential and integral equations with stochastic simulations. While we applied these methodologies specifically to the example at hand, we expect that combinations of such approaches can be relevant in wider context of gene-expression and biological modelling.

## Acknowledgements

PB is supported by the Slovak Research and Development Agency under the contract No. APVV-14-0378 and also by the VEGA grant 1/0347/18.

## References

[1] M.B. Elowitz, A.J. Levine, E.D. Siggia, and P.S. Swain, Stochastic gene expression in a single cell, Science 297 (2002), 1183–1186.

[2] D.R. Larson, R.H. Singer, and D. Zenklusen, A single molecule view of gene expression, Trends Cell Biol. 19 (2009), 630–7.

[3] L. Cai, N. Friedman, and X.S. Xie, Stochastic protein expression in individual cells at the single molecule level, Nature 440 (2006), 358–362.

[4] R. D. Dar, B. S. Razooky, A. Singh, T. V. Trimeloni, J. M. McCollum, C. D. Cox, M. L. Simpson, and L. S. Weinberger, Transcriptional burst frequency and burst size are equally modulated across the human genome, P. Natl. Acad. Sci. USA 109 (2012), 17454–17459.

[5] U. Alon, An Introduction to Systems Biology: Design Principles of Biological Circuits, Chapman & Hall/CRC, 2007.

[6] N. Friedman, L. Cai, and X.S. Xie, Linking stochastic dynamics to population distribution: an analytical framework of gene expression, Phys. Rev. Lett. 97 (2006), 168302.

[7] Miquel Angel Schikora-Tamarit, Carlos Toscano-Ochoa, Julia Domingo Espinos, Lorena Espinar, and Lucas B Carey, A synthetic gene circuit for measuring autoregulatory feedback control, Integr. Biol. 8 (2016), 546–555.

[8] Pavol Bokes and Abhyudai Singh, Gene expression noise is affected differentially by feedback in burst frequency and burst size, J. Math. Biol. 74 (2017), 1483–1509.

[9] P. Bokes, Y.T. Lin, and A. Singh, High cooperativity in negative feedback can amplify noisy gene expression, B. Math. Biol. (2018), doi: 10.1007/s11538–018–0438–y.

[10] J. Peccoud and B. Ycart, Markovian modeling of gene-product synthesis, Theor. Popul. Biol. 48 (1995), 222–234.

[11] Somkid Intep and Desmond J Higham, Zero, one and two-switch models of gene regulation, Discrete Cont. Dyn-B. 14 (2010), 495–513.

[12] Genghong Lin, Jianshe Yu, Zhan Zhou, Qiwen Sun, and Feng Jiao, Fluctuations of mrna distributions in multiple pathway activated transcription, Discrete Cont. Dyn-B. (2018). DOI: 10.3934/dcdsb.2018219.

[13] Renaud Dessalles, Vincent Fromion, and Philippe Robert, A stochastic analysis of autoregulation of gene expression, J. Math. Biol. (2017). DOI: 10.1007/s00285-017-1116-7.

[14] Peter Czuppon and Peter Pfaffelhuber, Limits of noise for autoregulated gene expression, J. Math. Biol. (2018). DOI: 10.1007/s00285-018-1248-4.

[15] Chen Jia, Hong Qian, Min Chen, and Michael Q Zhang, Relaxation rates of gene expression kinetics reveal the feedback signs of autoregulatory gene networks, J. Chem. Phys. 148 (2018), 095102.

[16] Haohua Wang, Peijiang Liu, Qingqing Li, and Tianshou Zhou, Entangled signal pathways can both control expression stability and induce stochastic focusing, FEBS Lett. 592 (2018), 1135–1149.

[17] J. Ren, F. Jiao, Q. Sun, M. Tang, and J. Yu, The dynamics of gene transcription in random environments, Discrete Cont. Dyn-B. (2018). DOI: 10.3934/dcdsb.2018224.

[18] Guilherme CP Innocentini, Michael Forger, Ovidiu Radulescu, and Fernando Antoneli, Protein synthesis driven by dynamical stochastic transcription, B. Math. Biol. 78 (2016), 110–131.

[19] Frits Veerman, Carsten Marr, and Nikola Popović, Time-dependent propagators for stochastic models of gene expression: an analytical method, J. Math. Biol. 77 (2018), 261–312.

[20] Guilherme CP Innocentini, Sarah Guiziou, Jerome Bonnet, and Ovidiu Radulescu, Analytic framework for a stochastic binary biological switch, Phys. Rev. E 94 (2016), no. 6, 062413.

[21] N. Kumar, T. Platini, and R. V. Kulkarni, Exact distributions for stochastic gene expression models with bursting and feedback, Phys. Rev. Lett. 113 (2014), 268105.

[22] A. Crudu, A. Debussche, A. Muller, O. Radulescu, et al., Convergence of stochastic gene networks to hybrid piecewise deterministic processes, Ann. Appl. Probab. 22 (2012), 1822–1859.

[23] Jakub Jedrak and Anna Ochab-Marcinek, Time-dependent solutions for a stochastic model of gene expression with molecule production in the form of a compound poisson process, Phys. Rev. E 94 (2016), 032401.

[24] P. Bokes, J.R. King, A.T.A. Wood, and M. Loose, Transcriptional bursting diversifies the behaviour of a toggle switch: hybrid simulation of stochastic gene expression, B. Math. Biol. 75 (2013), 351–371.

[25] Yen Ting Lin and Nicolas E Buchler, Efficient analysis of stochastic gene dynamics in the non-adiabatic regime using piecewise deterministic markov processes, J. Roy. Soc. Interface 15 (2018), 20170804.

[26] Yen Ting Lin, Peter G Hufton, Esther J Lee, and Davit A Potoyan, A stochastic and dynamical view of pluripotency in mouse embryonic stem cells, PloS Comput. Biol. 14 (2018), e1006000.

[27] Stefanie Winkelmann and Christof Schütte, Hybrid models for chemical reaction networks: Multiscale theory and application to gene regulatory systems, J. Chem. Phys. 147 (2017), 114115.

[28] Manuel Pájaro, Antonio A Alonso, Irene Otero-Muras, and C Vázquez, Stochastic modeling and numerical simulation of gene regulatory networks with protein bursting, J. Theor. Biol. 421 (2017), 51–70.

[29] Ryszard Rudnicki and Marta Tyran-Kamińska, Piecewise deterministic processes in biological models, Vol. 1, Springer, 2017.

[30] A. Raj, C.S. Peskin, D. Tranchina, D.Y. Vargas, and S. Tyagi, Stochastic mRNA synthesis in mammalian cells, PLoS Biol. 4 (2006), e309.

[31] Y. T. Lin and C. R. Doering, Gene expression dynamics with stochastic bursts: Construction and exact results for a coarse-grained model, Phys. Rev. E 93 (2016), 022409.

[32] Michio Masujima, Applied mathematical methods in theoretical physics, John Wiley & Sons, 2009.

[33] M.W. Hirsch, S. Smale, and R.L. Devaney, Differential Equations, Dynamical Systems, and an Introduction to Chaos, Academic Press, 2004.

[34] M. Abramowitz and I.A. Stegun, Handbook of Mathematical Functions with Formulas, Graphs, and Mathematical Tables, National Bureau of Standards, Washington, D.C., 1972.

